# Environmental DNA metabarcoding of rivers: Not all eDNA is everywhere, and not all the time

**DOI:** 10.1101/164046

**Authors:** Jan-Niklas Macher, Florian Leese

## Abstract

Environmental DNA metabarcoding has become a popular tool for the assessment of freshwater biodiversity, but it is largely unclear how sampling time and location influence the assessment of communities. Abiotic factors in rivers can change on small spatial and temporal scale and might greatly influence eDNA metabarcoding results. In this study, we sampled three German rivers at four locations per sampling site: 1. Left river side, surface water 2. Right river side, surface water, 3. Left side, close to the riverbed, 4. Right side, close to the riverbed. For the rivers Ruhr and Möhne, sampling was conducted three times in spring, each sampling one week apart. The Ruhr was again sampled in autumn and the Gillbach was sampled in winter. Sequencing on an Illumina MiSeq with COI primers Bf2/BR2 revealed diverse communities (6493 Operational taxonomic units, OTUs), which largely differed between rivers. Communities changed significantly over time in the Ruhr, but not in the Möhne. Sampling location influenced recovered communities in the Möhne and in the Ruhr in autumn. Our results have important implications for future eDNA studies, which should take into account that not all eDNA in rivers is everywhere and not at all times.

## Introduction

Worldwide, a threatening loss of biodiversity and ecosystem functions connected to rivers is observed and large-scale assessments of biodiversity are urgently needed to monitor this loss^1^. For that purpose, environmental DNA metabarcoding is especially promising. Environmental DNA (eDNA) metabarcoding means that DNA is extracted from soil, air or water and millions of sequences of a genetic marker are amplified and sequenced, allowing to assess a large part of an ecosystem’s biodiversity^2^. Recent findings show that environmental DNA metabarcoding can be used to assess whole river catchment biodiversity^3^, which is especially promising for large-scale studies. At the moment, freshwater monitoring programs largely rely on the sampling and morphological identification of invertebrate taxa, which can be error-prone^4^ and miss out on ecological differences between closely related species^5^. While whole community assessments of freshwater ecosystems using environmental DNA metabarcoding are still not common, they have been shown to be a potentially powerful tool^3,6,7^, which might greatly improve speed and accuracy of biodiversity assessments. However, the technique suffers from several potential drawbacks, which might hamper its usability. While tests have been conducted on the power of eDNA metabarcoding for the detection of species^8^, degradation speed of environmental DNA^9^, filtration^10^ and DNA extraction methods^11^, temporal patterns of eDNA occurrence^6^, very few studies have looked at spatial distribution of environmental DNA, especially in rivers. Studies have assessed communities recovered from eDNA samples in different stream habitats^12^, have shown that sediments function as a depots for DNA^13^ and that transport of eDNA in rivers depends on several factor such as retention and resuspension^14^. To date, however, many questions regarding eDNA transport, spatial resolution and reliability of results are still not resolved^15^. The question if and how eDNA is distributed in the water column of rivers on a very small scale has not been addressed. It is often assumed that at least in shallow rivers, most eDNA is everywhere due to the relatively turbulent water movement, which will disperse DNA in the water. However, rivers are very heterogeneous environments with a multitude of microhabitats, which can harbour highly different species communities on a small spatial scale^16^. If eDNA is to be used in biodiversity assessments of rivers, it needs to be clarified whether all eDNA is indeed everywhere at all times and if it is homogeneously distributed in the water column. If this is the case, sampling would be greatly simplified, as taking a water sample at any point in the stream would be sufficient for reliable biodiversity assessments. If differences on small spatial and temporal scale are found, however, this needs to be taken into account in future studies using environmental DNA, as even slightly shifted sampling times and locations could lead to different assessment results, subsequently biasing conclusions based on these findings.

In this study, we assessed the impact of sampling time and sampling location at a given sampling site on community composition recovered through eDNA metabarcoding. We sampled three rivers at four locations each and sequenced a 421 bp region of the standard animal barcoding gene COI using highly degenerate primers. We sampled surface water on the left and right side of the rivers and also took water samples directly beneath these surface sampling locations, close to the riverbed. Two rivers were sampled three times in spring, with each sampling one week apart, one of these rivers was sampled a fourth time five month later and one river was sampled once in winter. We hypothesise

1. that community composition inferred through eDNA metabarcoding strongly differs between rivers, even those of the same stream type and and in close proximity to each other
2. that community composition inferred through eDNA metabarcoding changes over the course of one week in rivers and
3. that community composition inferred from eDNA metabarcoding differs between different locations at a sampling site on a small scale, meaning that environmental DNA is not evenly distributed in the water columns of rivers.

## Results

### Discovered biodiversity

26,142,204 raw reads were obtained (MiSeq run 1: 16,005,709 MiSeq run 1, MiSeq run 2: 10,136,495). A total of 8,939,194 reads remained after initial quality filtering and 14,127 OTUs were retained. 1607 reads (0.018 % of reads) were found in negative control samples. This a commonly found read number due to tag switching that occurs during Illumina sequencing and thus, no contamination was suspected. After further quality filtering, i.e. retaining only OTUs present in both replicates per sample, 6493 OTUs (47.8%) were retained (read number per replicate: Supplementary table 1). On kingdom level (NCBI taxonomy), 39% of all OTUs from the Gillbach could be assigned to a taxonomic name, 43% of OTUs from the Möhne, 39% of OTUs from the Ruhr spring samples and 47% of the OTUs found in the Ruhr autumn samples. A large number of identified OTUs were Stramenopiles (Gillbach: 42%, Figure 1 a; Möhne: 54%, Figure 1 b; Ruhr (spring): 49%, Figure 1 c; Ruhr (autumn): 52%, Figure 1 d). Metazoa were the second largest group with 16% of OTUs in the Gillbach, 18% in the Möhne, 17% in the Ruhr in spring and 17% in the Ruhr in autumn. (Detailed list: Supplementary table 2). When using a strict threshold (~97% identity to reference sequences), 155 metazoan OTUs and 67 Stramenopiles were retained for further analyses (full OTU table: Supplementary table 3). In total, 42 metazoan orders were found. Diptera contributed most of the metazoan OTUs (21.3%), followed by Haplotaxida (11.6%), and Trichoptera (9.7%) (detailed list: Supplementary table 4). The identified Stramenopiles fell into 15 orders, with the majority of taxa being Oomycota (31.3 % Pythiales, 23.9% Peronosporales) (detailed list: Supplementary table 5).

**Table 1:** Results of PERMANOVAs (R^2^ effect sizes and significance) inferred through eDNA metabarcoding of the rivers Ruhr, Möhne and Gillbach for all OTUs, metazoan OTUs and stramenopile OTUs.

| Dataset | Differences between communities in Ruhr, Möhne and Gillbach |
| --- | --- |
| All OTUs | $R^2 = 0.55$, $p = 0.001$ |
| Metazoa | $R^2 = 0.43$, $p = 0.001$ |
| Stramenopiles | $R^2 = 0.56$, $p = 0.001$ |

**Table 2:** Results of the PERMANOVAs (R^2^ effect sizes and significance) describing the impact of sampling time on community composition in the Ruhr (spring + autumn; spring only) and the Möhne datasets for all OTUs, Metazoa and Stramenopiles.

| All OTUs | Differences between communities found in spring week 1, 2 and 3 |
| --- | --- |
| Ruhr Complete | $R^2=0.35$, $p=0.001$ |
| Ruhr: spring | $R^2=0.31$, $p=0.003$ |
| Möhne: spring | $R^2=0.08$, $p=0.831$ |
| Metazoa |  |
| Ruhr Complete | $R^2=0.26$, $p=0.001$ |
| Ruhr: spring | $R^2=0.39$, $p=0.001$ |
| Möhne: spring | $R^2=0.07$, $p=0.72$ |
| Stramenopiles |  |
| Ruhr Complete | $R^2=0.34$, $p=0.001$ |
| Ruhr: spring | $R^2=0.27$, $p=0.031$ |
| Möhne: spring | $R^2=0.11$, $p=0.339$ |

**Table 3:** Results of the PERMANOVAs (R^2^ effect sizes and significance) describing the impact of sampling location on community composition in the Ruhr (spring + autumn; spring only; autumn only) and the Möhne datasets for all OTUs, Metazoa and Stramenopiles.

| All OTUs | Sampling location | Surface vs. riverbed | Left side vs. right side |
| --- | --- | --- | --- |
| **Ruhr Complete** | $R^2=0.06$, $p=0.983$ | $R^2=0.02$, $p=0.824$ | $R^2=0.07$, $p=0.181$ |
| **Ruhr: spring** | $R^2=0.17$, $p=0.956$ | $R^2=0.05$, $p=0.949$ | $R^2=0.06$, $p=0.604$ |
| **Ruhr: autumn** | $R^2=0.31$, $p=0.12$ | $R^2=0.13$, $p=0.039$ | $R^2=0.09$, $p=0.411$ |
| **Möhne:spring** | $R^2=0.36$, $p=0.003$ | $R^2=0.11$, $p=0.127$ | $R^2=0.11$, $p=0.138$ |
| **Metazoa** |  |  |  |
| **Ruhr Complete** | $R^2=0.06$, $p=0.984$ | $R^2=0.02$, $p=0.891$ | $R^2=0.09$, $p=0.058$ |
| **Ruhr: spring** | $R^2=0.13$, $p=0.986$ | $R^2=0.03$, $p=0.951$ | $R^2=0.06$, $p=0.714$ |
| **Ruhr: autumn** | $R^2=0.35$, $p=0.041$ | $R^2=0.16$, $p=0.009$ | $R^2=0.11$, $p=0.218$ |
| **Möhne: spring** | $R^2=0.31$, $p=0.205$ | $R^2=0.18$, $p=0.013$ | $R^2=0.07$, $p=0.768$ |
| **Stramenopiles** |  |  |  |
| **Ruhr Complete** | $R^2=0.06$, $p=0.912$ | $R^2=0.03$, $p=0.533$ | $R^2=0.07$, $p=0.169$ |
| **Ruhr: spring** | $R^2=0.24$, $p=0.604$ | $R^2=0.14$, $p=0.179$ | $R^2=0.05$, $p=0.677$ |
| **Ruhr: autumn** | $R^2=0.28$, $p=0.434$ | $R^2=0.11$, $p=0.285$ | $R^2=0.1$, $p=0.371$ |
| **Möhne: spring** | $R^2=0.33$ 0.281 | $R^2=0.1$ 0.379 | $R^2=0.07$, $p=0.542$ |

**Figure 1:**
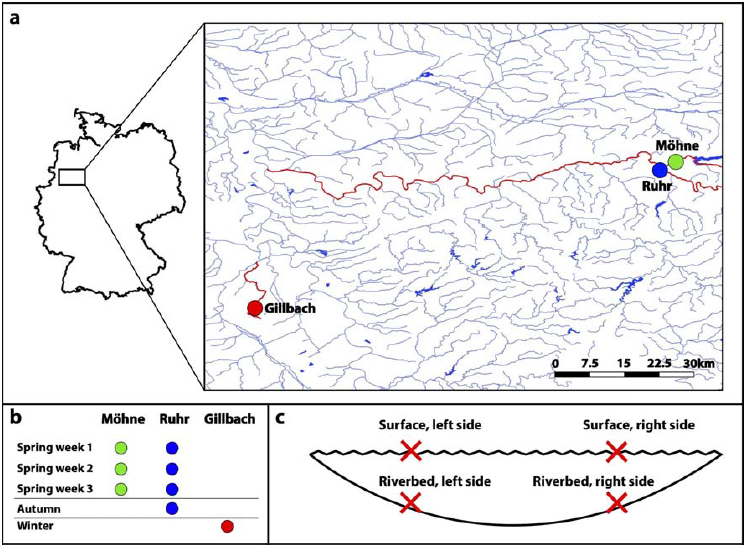
Taxonomy assigned to OTUs on kingdom level (NCBI taxonomy). a) Gillbach, b) Möhne, c) Ruhr (spring), d) Ruhr (autumn)

### Differences in community composition between rivers

Results for the datasets used for comparing community composition in different rivers can be found in table 1. Here, only PERMANOVA results with R^2^ >0.09 and p<0.05 are further described. Sampled rivers strongly differed with regards to community composition (All OTUs: R^2^=0.55, p=0.001; Metazoa: R^2^=0.43, p=0.001; Stramenopiles: R^2^=0.56, p=0.001), which is also shown by NMDS plots (All OTUs: Figure 2 a; Metazoa: Figure 2 b; Stramenopiles: Figure 2 c)

**Figure 2:**
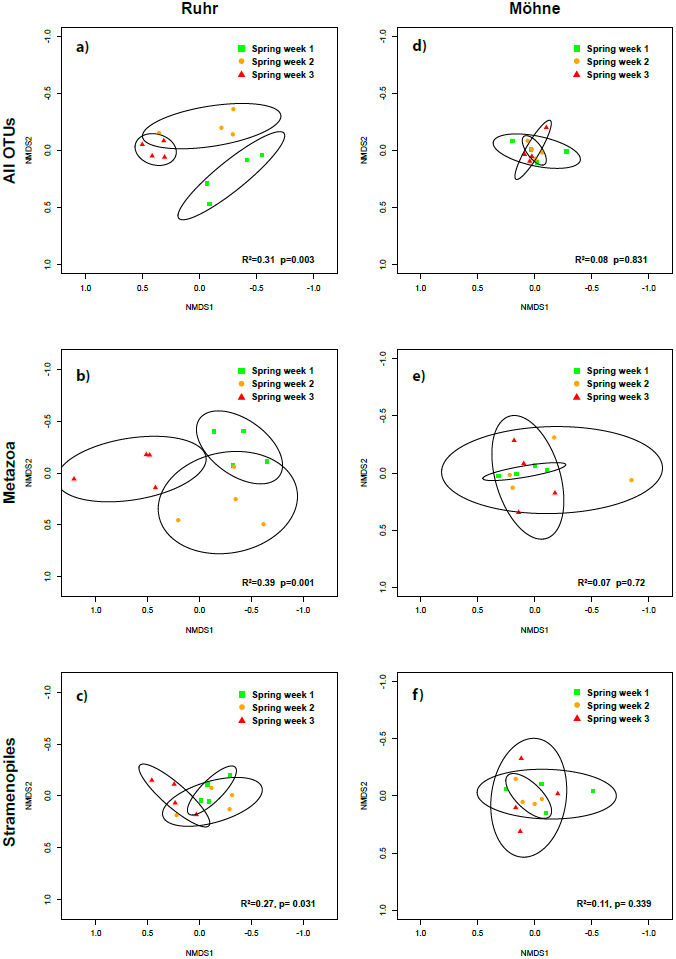
NMDS plot showing differences between communities in the rivers Ruhr (spring + autumn), Möhne and Gillbach. a) All OTUs, b) Metazoa, c) Stramenopiles. R^2^ and p values shown correspond to the respective PERMANOVA results.

### Influence of sampling time on communities inferred through eDNA metabarcoding

Results of all PERMANOVAs for the datasets comparing community composition at different sampling times can be found in table 2. Sampling time was found to strongly influence community composition of all OTUs (R^2^=0.35, p=0.001), Metazoa (R^2^=0.26, p=0.001) and Stramenopiles (R^2^=0.34, p=0.001) in the Ruhr (spring + autumn). The same was found when only the spring samples were analysed (All OTUs: R^2^=0.31, p=0.003, NMDS: Figure 3a; Metazoa: R^2^=0.39, p=0.001, NMDS: Figure 3b; Stramenopiles: R^2^=0.27, p=0.031, NMDS: Figure 3c). For the Möhne samples, which were taken at the same time as the Ruhr spring samples, sampling time did not significantly influence community composition (All OTUs: R^2^=0.08, p=0.831, NMDS: Figure 3d; Metazoa: R^2^=0.07, p=0.72, NMDS: Figure 3e; Stramenopiles: R^2^=0.11, p=0.339, NMDS: Figure 3f).

**Figure 3:**
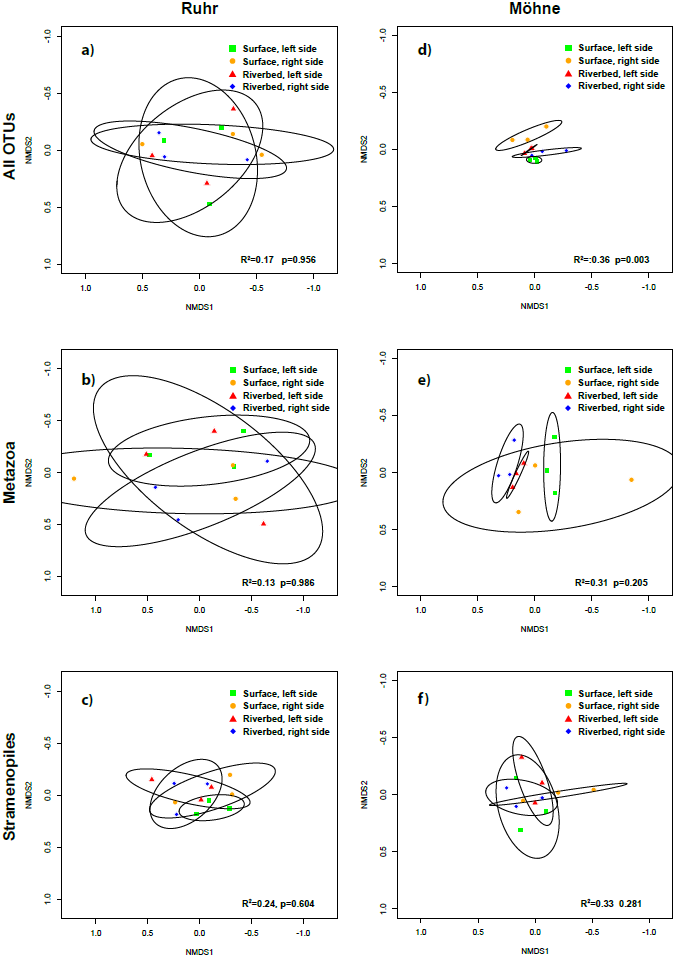
NMDS plot showing impact of sampling time (spring week 1, 2 and 3) on community composition in the rivers Ruhr and Möhne in spring. a) Ruhr, all OTUs, b) Ruhr, Metazoa, c) Ruhr, Stramenopiles, d) Möhne, all OTUs, e) Möhne, Metazoa, f) Möhne, Stramenopiles. R^2^ and p values shown correspond to the respective PERMANOVA results.

### Influence of sampling location on communities inferred through eDNA metabarcoding

All results of the PERMANOVAs for the datasets comparing community composition at different sampling locations can be found in table 3. Sampling location did not explain a significant proportion of the community composition in the Ruhr in spring (NMDS: Figure 4a, 4b, 4c) and autumn. Three samples were taken per sampling location in the Ruhr in autumn. Results show that these replicates do not always cluster closest together for all sampling locations (Supplementary Figure 1). However in autumn, community composition of all OTUs in the surface water of the Ruhr differed moderately from community composition found in the water sampled close to the riverbed (R^2^=0.13, p=0.039), which was also found for Metazoa (R^2^=0.16, p=0.009), but not for Stramenopiles (R^2^=0.11, p=0.285). Community composition in samples from left and right side of the Ruhr did not differ in autumn. For the Möhne samples, sampling location explained a large fraction of the variance in community composition of all OTUs (R^2^=0.36, p=0.003, NMDS: Figure 4d), but not that of Metazoa (R^2^=0.31, p=0.205, NMDS: Figure 4e) and Stramenopiles (R^2^=0.33, p=0.281, NMDS: Figure 4f). Community composition found in surface water of the Möhne differed from that found in riverbed water only for metazoan OTUs (R^2^=0.18, p=0.013). No significant differences were found between communities found in samples taken from the left and the right side of the stream.

**Figure 4:**
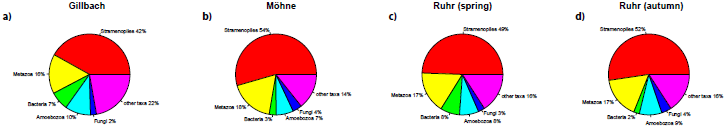
NMDS plot showing impact of sampling location on community composition in the rivers Ruhr and Möhne in spring. a) Ruhr, all OTUs, b) Ruhr, Metazoa, c) Ruhr, Stramenopiles, d) Möhne, all OTUs, e) Möhne, Metazoa. f) Möhne, Metazoa. R^2^ and p values shown correspond to the respective PERMANOVA results.

## Discussion

In this study, we investigated how time and sampling location influence community composition inferred through environmental DNA metabarcoding of rivers. We sampled three shallow rivers at four locations in close proximity to each other and two of these rivers were sampled at several points in time. We could show that not all environmental DNA in rivers is everywhere, and not all the time, even on a very small spatial scale.

We hypothesised that communities from different rivers, even those in close proximity to each other, significantly differ when inferred through eDNA metabarcoding. We could clearly show that this is true even for biocoenotically similar rivers like the Ruhr and Möhne, which are in close proximity (<10km) to each other. This finding shows the potential of the eDNA metabarcoding technique as a potential tool for ‘fingerprinting’ rivers and its potential to be used as a tool in routine monitoring, for which it has been proposed before^3^. Previous studies have shown that eDNA is capable of recovering differences in community composition between habitats^17,18^ and that communities in single locations change over time^6^. Due to the large amount of OTUs obtained (here: 6493), it can be expected that even minor changes in community composition of a river can be easily detected and used to quantify environmental changes of both natural (e.g. seasonal) and anthropogenic (e.g. chemical pollution) origin. However, the limited number of rivers included in our study does not allow to draw the definite conclusion that ‘fingerprinting’ rivers is possible, and further studies including a large number of rivers sampled at different points in time and at different sites are needed to answer the question if ‘stream fingerprinting’ by eDNA metabarcoding is possible and reliable.

Second, we hypothesised that community composition in rivers inferred through eDNA metabarcoding changes over time, as has been shown for invertebrates in lakes^6^ and fish in estuaries^18^. Although the fact that communities change over time due to biotic (presence of species) and abiotic factors (e.g. flow and water temperature) is known^19^, our study is the first to study this pattern in different rivers on a small temporal scale using eDNA and primers that target a wide range of taxa instead of selected taxonomic groups. We found different temporal patterns in the rivers Ruhr and Möhne: In the Ruhr, community composition of all OTUs, metazoan OTUs, and stramenopile OTUs changed from spring to autumn, which complements previous studies on community change driven by seasonality^6^. However, community composition in the Ruhr also significantly changed within two weeks in spring for all OTUs, metazoan OTUs and stramenopile OTUs. The latter pattern was not found for the Möhne, in which community composition remained similar during the same two weeks. We assume that changes in community composition in the Ruhr can be explained by minimal changes of abiotic factors, such as slightly prolonged duration of sunshine and higher temperatures or slight variations in discharge or flow velocity. Although similar in structure to the Möhne, the Ruhr river is widely unregulated above the sampling site. The Möhne, in contrast, is regulated by a dam approximately 10 kilometres upstream of the sampling site, which prevents stronger changes in discharge and flow velocity. This might explain the greater stability of community composition observed in the Möhne, while the community in the Ruhr might be more exposed to changes in the environment. Further studies over a longer period of time, including more rivers and measuring more abiotic factors on a finer scale are needed to definitely answer the question of which and how abiotic factors influence the community inferred through eDNA metabarcoding at different points in time. Independent of the reasons behind the observed patterns, our findings show that communities inferred through eDNA metabarcoding can change within a relative short time period. This has previously been found for biofilms^20^ and bacterioplancton^21^ in rivers, but to our knowledge, our study is the first to report that this pattern can also be observed for metazoan taxa when using environmental DNA metabarcoding. Our results have important implications for study design and sampling campaigns implementing eDNA, as they show that eDNA samples for studies should be taken in as little time as possible to prevent possible biases due to temporal variation.

Third, we hypothesised that community composition inferred from eDNA metabarcoding differs between different locations at a sampling site on a small spatial scale, meaning that environmental DNA is not evenly distributed in the water columns of shallow rivers. This has not been studied before, although it is known from non-molecular work that community composition in rivers changes with water depth^22^ and eDNA studies have found sediment to hold more fish DNA than surface water^13^, possibly hinting at a spatial pattern of eDNA distribution in rivers. We found contrasting patterns: In the Ruhr, sampling location did not explain community composition in spring, but in autumn when metazoan OTUs were analysed. However, our results also show that the three samples taken per sampling location in the Ruhr in autumn do not always cluster closest together. This highlights that community composition inferred through eDNA metabarcoding can change even when samples are taken within less than a minute, but it might also show that during processing and amplification of samples, community composition can be changed due to stochasticity of PCR reactions^23^. In the Möhne, sampling location explained the community composition of all OTUs. Likewise, communities found in surface water samples and those found in riverbed samples significantly differed in the Ruhr in autumn when analysing all OTUs and metazoan OTUs and also differed in the Möhne when metazoan OTUs were analysed.

Our finding shows that not all eDNA is homogeneously distributed in the water column in all rivers. In addition, as a spatial variation in community composition was found in the Ruhr in autumn, but not in spring, it shows that this pattern might change within a river over time, although statistical power was better for the Ruhr autumn samples due to the higher number of samples taken. It seems likely that in areas of high spring discharge due to precipitation or snowmelt, community composition might be more dynamic during these times, as has been shown before^24,25^. The same could happen at any time when events change abiotic factors in the stream. While horizontal sampling location did not significantly influence the inferred community composition, the vertical sampling location did. Communities change with water depth, e.g. due to different habitats^26,27^ and fish DNA has been shown to be more abundant in sediments^13^. However, environmental DNA metabarcoding has so far not been used to describe differences in community composition between surface and riverbed in rivers. Our finding that community composition inferred through eDNA metabarcoding can be different on a small spatial scale in shallow rivers has highly important implications for future eDNA-based work. If this pattern is found in shallow rivers with less than 1 metre depth, it can be expected to be even more pronounced in wide, deep rivers, which are known to harbour highly diverse communities on a small spatial scale^28^. Thus, if the goal is to assess the whole biodiversity, several samples of surface and riverbed water should be taken, an approach that is similar to the multi-habitat sampling used for classical sampling of invertebrates^29^ and that is also applied when sampling eDNA in standing waters^30,31^. Surface water might more often contain eDNA from further upstream in the river catchment, while riverbed water flows over a multitude of obstacles and might contain more eDNA from the (micro)habitats directly upstream of the sampling site. It has been shown previously that eDNA can be transported over long distances^32^, but also that retention and resuspension can play a major role^14^. To date it remains unclear to what extent transport distance of eDNA differs between surface water and water moving close to the riverbed and spatial resolution of eDNA is also not yet fully understood^15^. Our study highlights the need for follow-up studies addressing the community composition on small spatial scale. Sampling several upstream sites simultaneously might reveal where eDNA found at different locations in the water column of rivers actually originates.

Another aspect to take into consideration when planning eDNA studies is the choice of primers. We found the highly degenerate BF/BR2 primers, which were originally developed for stream macroinvertebrates^33^, to amplify a wide range of taxonomic groups. However, only between 16 % and 18 % of OTUs per stream were found to be Metazoa, which is probably due to the comparably large biomass of floating microorganisms in the water. The majority of OTUs (between 42 % and 54 % per stream) were identified as Stramenopiles, which comprise Oomycota and Bacillariophyta (diatoms). The COI gene can be used to identify taxa within these groups^34,35^ and other microbial taxa such as Amoeba^36^, but reference databases are currently poorly equipped, making taxonomic assignment challenging or impossible. For amplifying macrozoobenthic taxa relevant for classical stream monitoring, the BF2/BR2 primers have been shown to perform very well^37^, but more specific primers might be be a better choice when these taxa are to be amplified from eDNA water samples. Recently, long-range PCR with specific 16S rRNA primers has been shown to amplify whole mitochondrial genomes from eDNA samples, which might be an alternative for future eDNA metabarcoding studies when paired with high-throughput sequencing approaches on NGS machines producing long reads^38^. However, if assessing the whole community, including microbial taxa, is aimed for, using degenerate COI primers might be an alternative to current approaches using markers such as 18s rRNA, which is commonly used^39,40^. Although COI is not without problems when it comes to non-metazoan taxa^41^, mostly because little is known on how well the marker resolves taxa, using highly degenerate COI primers might be a promising approach to bridge the gap between metabarcoding microbial taxa and metazoans, for which mainly COI databases exist^42^.

In conclusion, we found strong evidence that not all eDNA is everywhere and not all the time in rivers. Future studies applying the technique of eDNA metabarcoding should be carefully planned with regard to sampling design and primer choice. It should be made sure to collect water for analyses within a short period of time and from several sampling locations in a river in order to prevent misinterpretations due to changing abiotic factors or different sampled microhabitats. In order to make the technique of eDNA metabarcoding widely applicable for biomonitoring and community ecology, more studies addressing the exact spatial and temporal scale of eDNA distribution and the impacts of abiotic factors on the recovered community are needed. This is especially important as eDNA is becoming an increasingly popular tool for biomonitoring and biodiversity assessment of ecosystems. The technique holds great potential, but technical issues such as appropriate primers and sampling design have not been widely tested, a step that should not be neglected if the technique is to be used in biodiversity assessments and ecological studies.

## Methods

Samples were taken from three rivers: Ruhr (51°26'55.6"N 7°57'09.0"E), Möhne (51°28'03.7"N 7°58'59.6"E) and Gillbach (51°00'52.2"N 6°41' 04.4"E) (Figure 5a). Ruhr and Möhne are similar in their characteristics in close proximity to each other (<10km). Ruhr and Möhne were sampled three times in spring, with each sampling one week apart. The Ruhr was again sampled in autumn and the Gillbach was sampled in winter (Figure 5b). Water samples were taken by filling a new, sterile plastic bottle (Nalgene, Rochester, USA) with 1 litre of water directly below the stream surface for the two surface sampling locations and directly below these locations, 5 cm above the riverbed for the riverbed sampling locations (Figure 5c). Stream depth was roughly 75 cm (Ruhr), 60cm (Möhne) and 45 cm (Gillbach).

**Figure 5:**
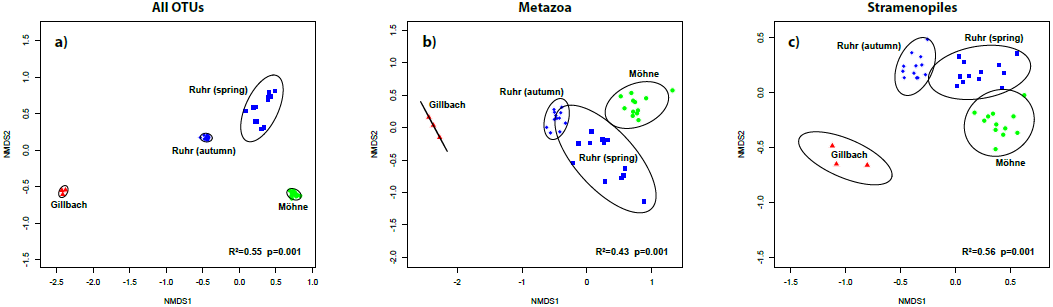
a) Map of the sampling sites, with sampled rivers marked in green (Möhne), blue (Ruhr) and red (Gillbach) b) Sampling scheme in spring, autumn and winter for the three sampled rivers. c) Profile of a stream showing sampling locations at a given site.

To test for variation of individual samples per location, three bottles of water per location were taken from the Ruhr in autumn. For taking the riverbed water samples, bottles were immersed, opened, filled and closed again under water. Sampling sites were marked with GPS and poles in the water to exactly sample the same locations during the next visits. Sampling locations were located roughly 1/4 of the stream width away from the left and right riverbanks, respectively (Möhne: 3.5 m, Ruhr: 8 m, Gillbach: 1.5 m). All samplings were performed between 11:00 am and 1 pm. Water temperature at each location was measured with a thermometer directly below the surface. Water depth was measured by using a folding rule and flow velocity was measured by letting a styrofoam ball drift over the distance of one meter while stopping the time (all abiotic data: Supplementary table 6). Bottles were transported back to the lab at 4 °C and 1 l of water was filtered through cellulose nitrate filters (0.2 μm pore size, Nalgene) with the help of a vacuum pump (VCP-8101). For each sampling, one litre of sterile water was filtered as negative control (eight total). Filters were carefully handled with sterile tweezers, folded, transferred to 90% molecular grade EtOH and stored at −20 °C until further processing. All lab work was conducted in a lab room specifically prepared and only used for eDNA work, in which no PCR products are present. All persons involved in handling samples wore full body protective clothing and surgical masks as breathing protection. The lab room is frequently cleaned with hydrogen peroxide solution and irradiated with high-intense UV light every night. Furthermore, samples were only handled under sterile UV hoods.

Filters were ripped to small pieces before proteinase k digestion and DNA extraction followed a salt extraction protocol (^43^ adjusted as in ^44^). Extracted DNA was quantified on a FragmentAnalyzer with the Standard Sensitivity Genomic Kit (AdvancedAnalytical, Oak Tree, USA) and 15 ng of DNA per sample was used for PCR. A two-step PCR protocol was used: For the first PCR, primers BF2 and BR2^33^ without tails were used for amplification using illustra PuReTaq Ready-to-go PCR beads (GE Heatltcare, Little Chalfont, UK). After an initial denaturation for 3 minutes at 94°C, 25 cycles at 94°C for 30 seconds, 48°C for 30 seconds, 72°C for two minutes were performed, followed by a final elongation at 72°C for five minutes. For the second PCR, 1 μl of the product was used with individually tagged BF2 / BR2 primers (combinations: Supplementary table 1). The PCR protocol remained the same, but 15 cycles were used. Two independent PCRs (1st + 2nd step) were run for each sample. PCR products were cleaned up using the MinElute PCR Purification Kit (Qiagen, Venlo, Netherlands), quantified on the Fragment Analyzer using the NGS High Sensitivity Kit, left side size selected using SpriSelect beads (Beckman Coulter, Brea, USA) and equimolar pooled. Negative controls were quantified together with the other samples and were added to the library so that they made up 10% of the total library volume. The final DNA library was again cleaned using the Qiagen MinElute kit and sent for sequencing on the Illumina MiSeq platform (Two runs, v2 chemistry, 2x250bp) at GATC Biotech (Constance, Germany).

Raw reads were processed as in^45^. In short, reads were demultiplexed and paired-end reads were merged using USEARCH (v.8.1.1756^46^). Since read quality of one sequencing run was low, read pairs were discarded if the number of expected errors predicted by the phred scores after merging was higher than three. Primers were removed with cutadapt (v.1.9^47^). Sequences were dereplicated, singletons were removed and the Uparse pipeline^48^ (97% identity) was used to cluster OTUs. Further, raeds including singletons were mapped against the clustered OTUs. Subsequently, only OTUs that had read abundances over 0.004 percent (i.e. read abundance >1 in the samples with =<25,000 reads) per sample were retained, while OTUs with lower read abundance were discarded (all scripts: Supplementary material 1). This is a suitable alternative to rarefaction^49^. Due to the higher threshold used for paired-end merging performed by USEARCH and in order to remove possible false OTUs, only OTUs present in both replicates per sample were counted as present and further analysed.

Taxonomy was assigned to OTUs using MEGAN^50^ (Settings: -evalue 1e-60, -max_target_seqs 10). To assign taxonomy to OTUs on kingdom level (NCBI taxonomy), a min_score of 300 (corresponding to ~80% identity in this dataset consisting of 421 bp reads was applied. Metazoan and stramenopile OTUs used for further analyses were identified with a restrictive min_score of 700 (corresponding to ~97% identity) to further filter out possible false OTUs. A custom made dataset consisting of all COI sequences deposited in Genbank (2.5 million sequences, downloaded on 20-02-2017, maximum length 5000 bp) was used for Megan analyses. Sequences were dereplicated prior to database building in order to remove overrepresented genetic sequences. The lowest taxonomic level assigned to OTUs was order to account for any inaccuracies due to misidentified specimens in the database.

Analyses of community structure were performed in R (v.3.3.2, R Core Development Team 2017) using the package vegan (v.2.4-1^51^) for all OTUs, metazoan OTUs and stramenopile OTUs. Jaccard distances were calculated with the vegdist function. The ‘adonis’ function was used to perform PERMANOVA as in^52^ on Jaccard distances calculated for communities in the three sampled rivers, per sampling location (surface left side vs. surface right side vs. riverbed left side vs. riverbed right side; left riverside vs. right river river side; surface vs. riverbed) and per sampling time, respectively, to test whether these factors explain community composition. The Gillbach was only included in analyses inferring the differences in community composition between rivers, since only three sampling locations were retained after read quality filtering.

Only PERMANOVA results with R^2^ >0.09 and p<0.05 were regarded as strong enough to reliably show an effect of the tested factor. Our approach is based on the assumption that a correlation coefficient r of 0.3 is showing a moderate effect^53^, and 0.09 is the corresponding R^2^. Consequently, we interpret R^2^ >0.25 as indicating strong effects, which corresponds to r >0.5 (a strong effect as defined by^54^). NMDS plots for the binary dataset were generated using the metaMDS function as implemented in vegan (binary =T, k = 2, trymax = 1000, autotransform = F). The abiotic data was used to visualise community structure with the ordiplot function. Maps were created with QGIS (v.2.18, QGIS Development Team, 2016) and figures were created with Adobe Illustrator (Adobe Systems, San José, USA)

## Data availability

Data has been deposited in the Short Read Archive (will be available upon publication)

## Acknowledgements

We thank Edith Vamos, Bianca Peinert and Cristina Hartmann-Fatu for help in the lab, Thomas Euteneuer-Macher and Anika Jedrzejewski for help with sampling and the Herzog-Sellenberg Foundation for financial support. Vasco Elbrecht, Romana Salis, Vera Ziska, Martina Weiss and Arne Beermann are thanked for proof-reading the manuscript and providing helpful comments.

## Author contributions

JNM and FL designed the experiment, JNM conducted field and laboratory work, JNM performed the analyses and JNM and FL wrote the manuscript.

